# AutoRNA: RNA tertiary structure prediction using variational autoencoder

**DOI:** 10.1101/2024.06.18.599511

**Authors:** M.A. Kazanskii, L. Uroshlev, F. Zatylkin, I. Pospelova, O. Kantidze, Y. Gankin

## Abstract

**Background:** Understanding the three-dimensional (3D) organization of RNA is essential for advancing therapeutic development and vaccine design. However, the limited availability of experimentally resolved RNA structures restricts the applicability of data-intensive machine learning approaches for tertiary structure prediction. This study aims to develop a data-efficient method for learning coarse-grained RNA structural organization directly from sequence information.

**Methods:** We propose AutoRNA, a variational autoencoder-based model that learns sequence-conditioned structural priors in the form of inter-nucleotide distance matrices. The model was trained on RNA structures obtained from the Protein Data Bank and restricted to sequences up to 64 nucleotides. Predicted distance matrices were converted into 3D coordinates using multidimensional scaling, followed by template-based assembly and molecular dynamics refinement. Model performance was evaluated using mean absolute error (MAE), root mean square error (RMSE), global distance test (GDT), and template modeling (TM) scores.

**Results:** On the test dataset, AutoRNA achieved an RMSE of 4.49 Å and an MAE of 3.13 Å for predicted inter-nucleotide centroid distances. However, the reconstructed three-dimensional structures showed limited global fold recovery, as indicated by moderate GDT scores and low TM-scores. Performance decreased for longer sequences, indicating limitations associated with data scarcity and increased structural complexity. Molecular dynamics refinement provided modest improvements, particularly for initially low-quality predictions.

**Conclusions:** AutoRNA demonstrates that variational autoencoders can learn meaningful coarse-grained structural representations of RNA from limited data. While not suitable for near-native tertiary structure prediction, the method generates candidate coarse-grained inter-nucleotide distance restraints that could potentially be incorporated into downstream structure-reconstruction or physics-based refinement workflows. This work highlights the potential of generative models for RNA structure prediction in low-data regimes.

## 1 Background

RNA molecules adopt complex three-dimensional (3D) structures that are closely linked to their biological functions, including regulation, catalysis, molecular recognition, and interactions with other biomolecules [1, 2]. Understanding RNA 3D structure is therefore important both for elucidating RNA function and for the development of RNA-targeted therapeutic strategies [2]. Leveraging machine learning (ML) techniques to decipher protein folding can solve challenges in the RNA domain. AlphaFold2 [3] (and the more recent AlphaFold3 [4]) have demonstrated the power of ML methods for solving protein-folding problems. Nevertheless, adapting neural networks similar to AlphaFold for RNA could present challenges.

The number and diversity of experimentally determined RNA structures available for model development remain substantially smaller than those available for proteins [5]. This relative scarcity of RNA structural data presents a challenge for data-intensive machine-learning approaches to RNA structure prediction. In addition, the availability and informativeness of evolutionary information derived from multiple-sequence alignments vary substantially among RNA families, motivating complementary approaches that can learn structural information directly from individual RNA sequences.

Evolutionary information obtained from multiple-sequence alignments can provide valuable constraints for RNA structure modeling. In particular, sequence covariation has been widely used to identify conserved base-pairing and structural relationships in RNA. However, the amount and statistical power of detectable covariation depend on the depth, diversity, and evolutionary variation of the available sequence alignments, and therefore vary among RNA families. Consequently, approaches that can learn structural information directly from individual RNA sequences may provide a complementary strategy, particularly when sufficiently informative multiple-sequence alignments are unavailable.

Here, we present a relatively simple variational autoencoder (VAE)-based architecture for learning sequence-conditioned coarse-grained structural priors of RNA molecules in the form of inter-nucleotide distance restraints. These predictions are intended to capture coarse-grained spatial organization rather than near-native atomic-level RNA structures. Our approach achieved an average root mean squared error (RMSE) of approximately 4.5 Å. In this work, we use the term “tertiary structure prediction” to refer to predicting coarse-grained 3D organization in the form of sequence-conditioned inter-nucleotide distance restraints, rather than recovering near-native atomic-level folds.

RNA 3D structure prediction has evolved from physics-based and coarse-grained sampling approaches toward hybrid and deep-learning methods. Early approaches included simplified lattice models [7] and coarse-grained Monte Carlo methods such as iFoldRNA [8]. Fragment-assembly and coarse-grained approaches remain relevant; for example, FARFAR2 combines fragment assembly, scoring, and high-resolution refinement [16], whereas modern implementations of SimRNA use coarse-grained conformational sampling with knowledge-based statistical potentials [48].

More recently, deep-learning approaches have substantially expanded the methodological landscape of RNA 3D structure prediction. trRosettaRNA predicts RNA structural restraints using a transformer-based architecture and combines these predictions with Rosetta-based structure construction [15]. RhoFold uses RNA language-model representations within a deep-learning framework for end-to-end RNA 3D structure prediction and has demonstrated improved accuracy on contemporary RNA structure-prediction benchmarks [33]. General biomolecular structure-prediction models, including AlphaFold3 [4] and Boltz-2 [32], further illustrate the increasing role of learned representations in modeling nucleic-acid structures and interactions. These recent developments provide the appropriate context for AutoRNA: rather than aiming at near-native end-to-end structure prediction, AutoRNA investigates whether a comparatively simple generative model trained on a limited structural dataset can learn sequence-conditioned coarse-grained inter-nucleotide distance restraints.

Statistical and knowledge-based potentials are widely used in RNA structure modeling to evaluate candidate three-dimensional conformations [9]. Molecular-mechanics force fields provide a different approach, in which molecular interactions are described using physically motivated energy terms and parameterized interaction potentials. Widely used biomolecular force-field families include AMBER [11] and CHARMM [12], which are commonly used in molecular dynamics simulations. Coarse-grained models [14] can reduce this computational complexity and enable more efficient conformational sampling. Hybrid methods have become increasingly popular in recent years. These methods combine empirical potentials with various neural network structures. For instance, one study applied the transformer architecture in conjunction with Rosetta’s empirical potentials [15]. Fragment-assembly methods provide another strategy for RNA tertiary structure prediction. For example, FARFAR2 performs de novo RNA structure prediction through fragment assembly followed by scoring and high-resolution refinement within the Rosetta framework [16]. Deep learning has also been successfully applied to RNA structural modeling at the secondary-structure level. For example, UFold uses a U-Net-based architecture to predict RNA secondary structures from sequence information [17].

RNA structure-prediction methods span a wide range of computational strategies, including physics-based simulation, fragment assembly, statistical potentials, and deep learning. Many of these approaches require either substantial conformational sampling, external structural information, or large-scale training data, depending on the method [14–16, 33]. In this study, we investigate whether a VAE trained on RNA distance matrices can learn sequence-conditioned coarse-grained structural representations from a relatively small structural dataset.

## 2 Methods

### 2.1 Data preprocessing and experiment

We used a PDB dataset containing the 3D structural information of RNA molecules [18]. We selected PDB structures containing RNA and excluded structures containing protein or DNA components. During preprocessing, duplicate entries and entries with empty RNA sequences were removed, and only RNA nucleotide residues were retained for subsequent structural analysis. For each nucleotide, we calculated a single representative three-dimensional coordinate as the geometric centroid of all atoms belonging to that nucleotide residue in the PDB structure. The centroid was calculated as

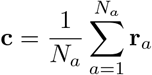

where *N*_*a*_ is the number of atoms in the nucleotide, **r**_*a*_ is the three-dimensional Cartesian coordinate of atom *a*, and **c** is the resulting nucleotide centroid. All atoms present in the corresponding nucleotide residue were included in the calculation. Thus, an RNA molecule containing *N* nucleotides was represented by *N* three-dimensional points, one per nucleotide. Pairwise Euclidean distances between these nucleotide centroids were then used to construct the distance matrix representing the RNA structure.

For the main experiment, we restricted the dataset to RNA sequences containing at most 64 nucleotides. This cutoff was selected because the available structural dataset was strongly enriched in short RNA sequences, while the number of available structures decreased substantially with increasing sequence length. In addition, the VAE uses a fixed-size 64 × 64 distance-matrix representation; therefore, restricting the primary analysis to sequences of at most 64 nucleotides provided a fixed input/output dimensionality while retaining the majority of the available short-RNA structures.

After preprocessing and application of the length cutoff, 852 RNA sequences were retained for the main experiment. A total of [X] otherwise eligible sequences exceeding 64 nucleotides were excluded by this cutoff. We generated a distance matrix by calculating the pairwise Euclidean distances between the nucleotide centroid coordinates. For nucleotide sequences containing fewer than 64 nucleotides, we applied zero padding.

We applied distance clipping at 100 Å to handle a very small number of extreme outliers. Quantitatively, the majority of pairwise distances fell within the range 0–80 Å, and only 0.06% of all distances exceeded 100 Å in the dataset containing sequences shorter than 64 nucleotides. Therefore, clipping affected only a negligible fraction of pairwise distances and was used to prevent rare extreme values from disproportionately influencing model training. For model input and training, pairwise distances were clipped at 100 Å and normalized to the range [0, 1] by division by 100:

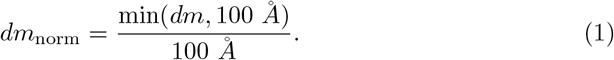

The multiplication of normalized values by 255 was performed only for grayscale visualization in Figure 1 and was not part of the model input or training procedure.

**Fig. 1.**
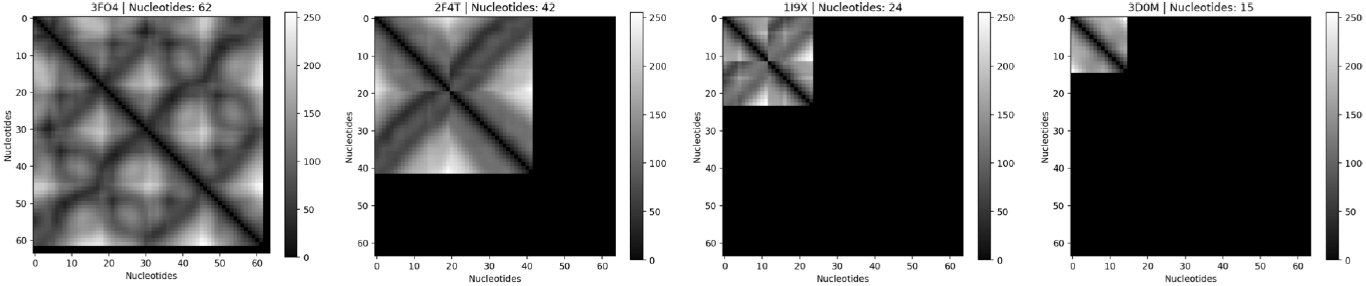
Examples of normalized RNA inter-nucleotide distance matrices with zero padding. Pairwise distances were clipped at 100 Å and normalized to the range [0, 1] by division by 100. For visualization only, the normalized values were multiplied by 255 to generate grayscale images. Nucleotides are represented along the x- and y-axes, and the black regions correspond to zero padding used to obtain a fixed matrix size of 64 *×* 64.

Together with the distance matrices of the RNA sequences, we used nucleotide letters A, G, C, and U as one-hot encoder vectors with the following rules:

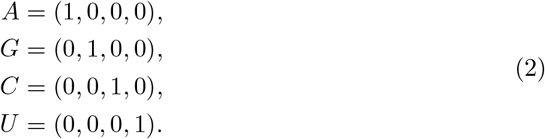

Where we added padding with values (0, 0, 0, 0) to the sequences of length less than 64.

To reduce sequence redundancy across the training, validation, and test subsets, pairwise sequence similarities were calculated using Biopython’s PairwiseAligner. For each sequence pair, the highest-scoring alignment was selected, and sequence similarity was calculated as the number of matching aligned positions divided by the length of the shorter sequence. A pairwise distance was then defined as 100−*S*, where *S* is the sequence-similarity percentage. Hierarchical agglomerative clustering was performed on the resulting distance matrix using average linkage, and clusters were obtained by cutting the hierarchy at a distance threshold of 60. The resulting clusters, rather than individual RNA structures, were assigned to the training, validation, and test subsets, such that all structures belonging to a given cluster remained within the same subset. This cutoff was selected as an empirical dataset-partitioning parameter rather than as a universal threshold for RNA homology or structural independence. We additionally examined a distance threshold of 70; this resulted in only 16 clusters, whereas a distance threshold of 60 produced 100 clusters. The larger number of clusters obtained with the threshold of 60 provided greater flexibility for constructing separate training, validation, and test subsets. The three largest clusters contained 111, 77, and 74 sequences, respectively.

Figure 2 visualizes the sequence clusters used to construct the training, validation, and test subsets. The visualization demonstrates the separation and relative organization of the sequence-similarity clusters obtained by hierarchical clustering. Principal component analysis (PCA) was used only to project the sequence-similarity relationships into two dimensions for visualization and was not used to define the clusters. Different colors indicate clusters identified by the hierarchical clustering procedure.

**Fig. 2.**
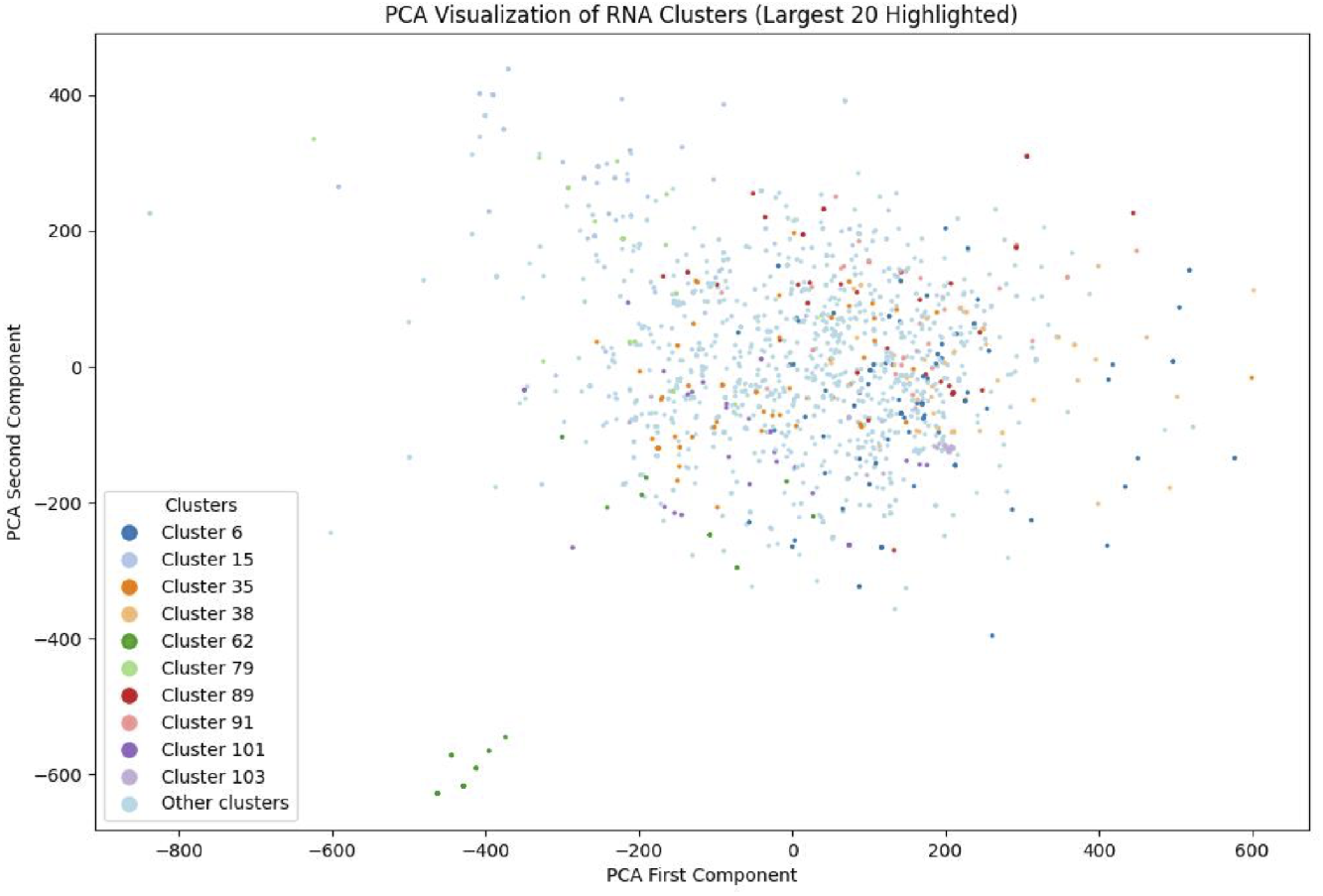
A graph of the first two principal components of the similarity matrix. Different colors represent distinct clusters obtained using hierarchical clustering.

We divided the dataset into training, validation, and testing subsets with approximate proportions of 80%, 15%, and 5%, respectively. A total of 852 sequences were obtained. After partitioning, there were 719, 96, and 37 RNA sequences in the training, validation, and testing subsets, respectively. Figure 3 presents the distribution of the lengths of the RNA sequences in the training, validation, and testing subsets and the total dataset.

**Fig. 3.**
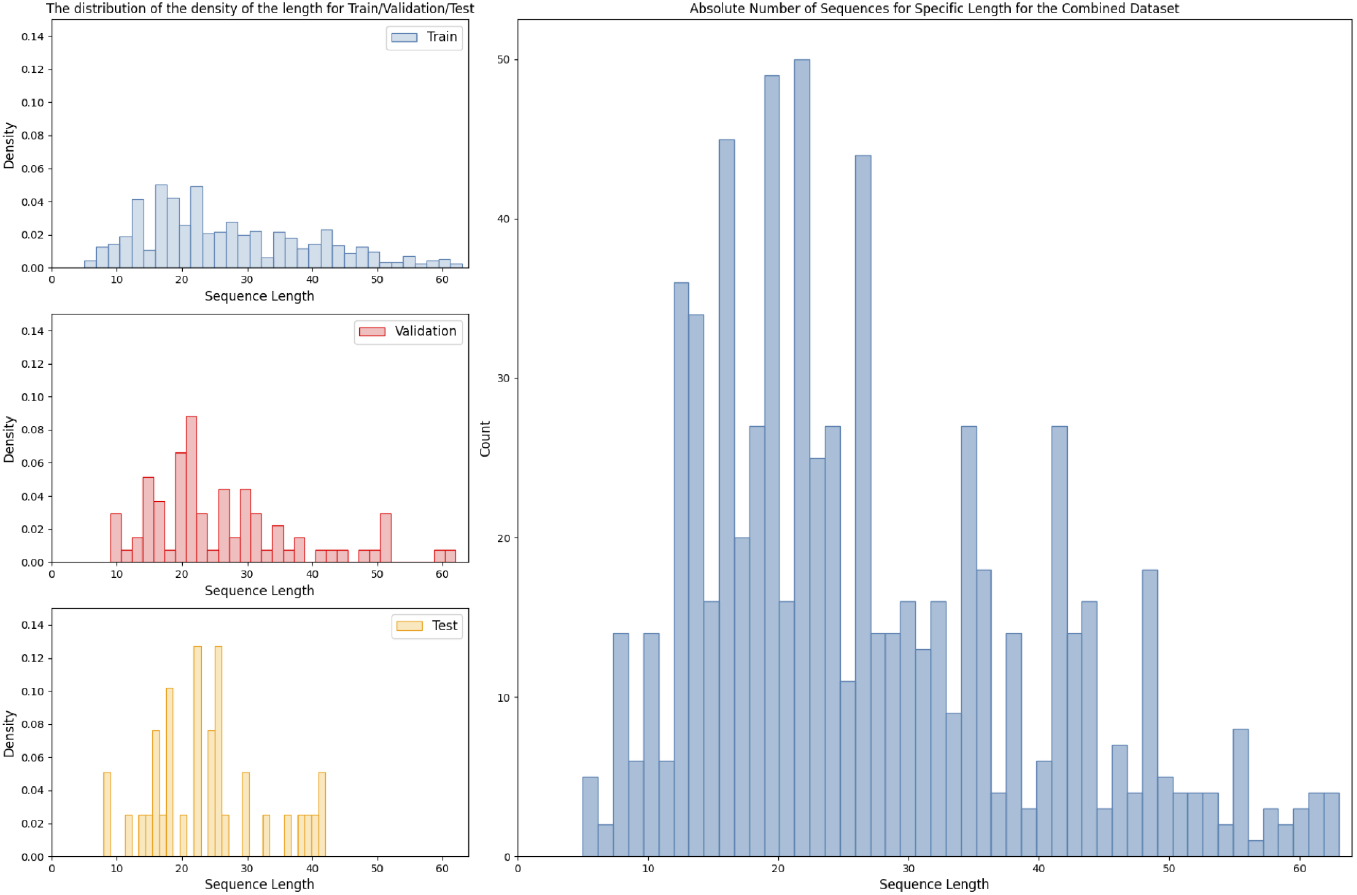
The probability density function of sequence lengths for the training, validation, and testing subsets (left) and the distribution of the sequence lengths in the combined dataset (right).

### 2.2 Pipeline

We used the VAE [22] to predict the distance matrix. The condition vector was the flattened one-hot encoded nucleotide sequence (256 elements). A stochastic input vector with the same dimensionality as the flattened distance matrix (64 × 64 = 4096 elements) was generated independently for each sample using values drawn from a standard normal distribution, *N*(0, 1). Python, NumPy, and PyTorch random-number generators were initialized using the experiment-specific random seed. We used a conditional variational autoencoder consisting of an encoder, a stochastic latent representation, and a decoder. The encoder contained fully connected hidden layers of 4096 and 2048 units. The encoder output was mapped to two separate 1024-dimensional vectors representing the latent mean (*µ*) and log-variance (log *σ*^2^). A 1024-dimensional latent vector was sampled using the standard VAE reparameterization procedure, *z* = *µ* + *σ* ⊙ *ϵ*, where *ϵ* ~ *N* (0, *I*). The decoder received the sampled latent vector together with the 256-dimensional sequence-condition vector and mapped this representation through fully connected hidden layers to a 4096-dimensional output representing the flattened 64 × 64 distance matrix. The last layer represented the values of the distance matrix. After each layer was fully connected (excluding the last one), we used batch normalization to enhance performance and stability. We also applied dropout at a rate of 0.5 for the output signals from the batch-normalization layers. Given the relatively small size of the available RNA structural dataset (852 samples) compared with the model capacity (approximately 64.5 million trainable parameters), dropout (rate = 0.5) was used as a stochastic regularization strategy intended to reduce the risk of overfitting. In addition, sequence-similarity-based clustering was applied prior to dataset partitioning to reduce sequence redundancy across the training, validation, and test subsets. This partitioning strategy reduced the likelihood that highly similar RNA sequences were assigned to different subsets; however, sequence-based clustering does not guarantee structural independence between the subsets.

The architecture of the VAE and the integration of the sequence condition with the model input are illustrated in Figure 4. Our choice of a fully connected VAE was motivated by the characteristics of the representation and the study objective. First, the input to the model is a global pairwise distance matrix, where every element depends on every other nucleotide; therefore, using dense layers ensures that no connectivity assumptions are imposed a priori, allowing the model to learn non-local tertiary dependencies directly from the data instead of constraining it to local receptive fields as in CNNs.

**Fig. 4.**
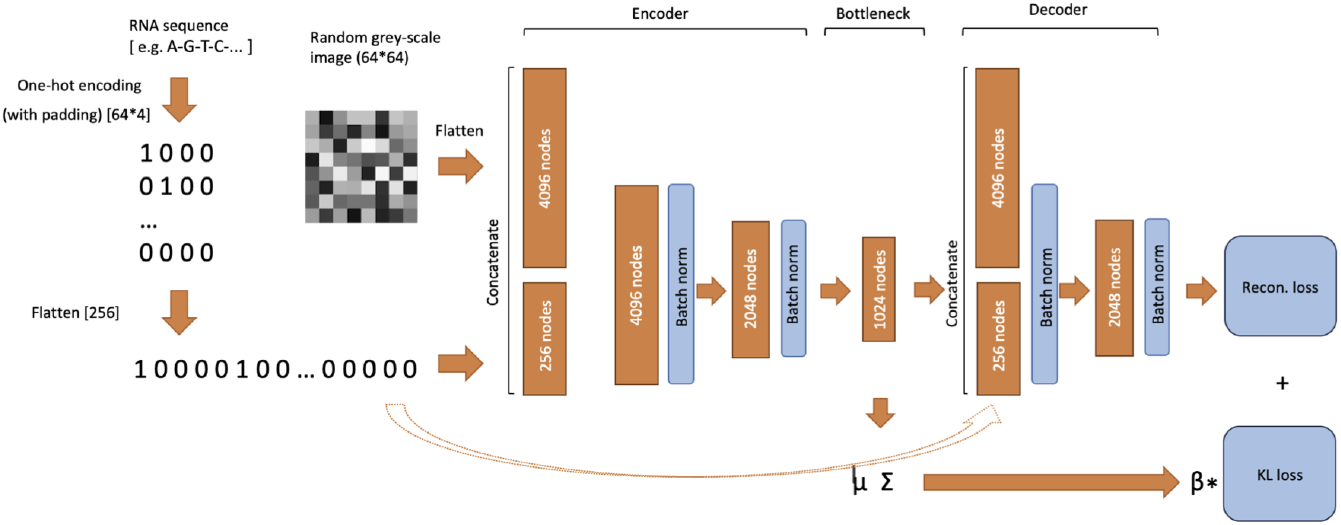
Schematic representation of the variational autoencoder (VAE). The RNA sequence is one-hot encoded and used as a conditioning vector. The encoder maps the input through fully connected hidden layers to separate latent mean (*µ*) and log-variance (log *σ*^2^) vectors. A 1024-dimensional latent representation is sampled using the VAE reparameterization procedure and passed to the decoder together with the sequence-condition vector. The decoder produces a 4096-dimensional output representing the flattened 64 *×* 64 inter-nucleotide distance matrix. The loss function combines the reconstruction loss based on mean absolute error (MAE) with the Kullback–Leibler (KL) divergence.

The model’s loss followed the standard formulation for variational autoencoders and consisted of two terms: the reconstruction loss and the Kullback–Leibler (KL) divergence [23]. The reconstruction term was computed using the mean absolute error (MAE), chosen for its robustness to outliers compared to the mean squared error (MSE). A weighting coefficient *β* was used to balance the reconstruction and regularization contributions. The total loss is

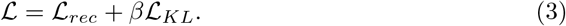

where the reconstruction loss is defined as

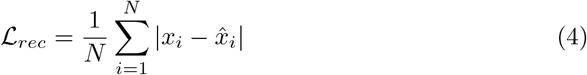

where *N* is the number of non-padding distance-matrix elements included in the loss calculation, *x*_*i*_ is the reference value, and 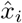 is the corresponding predicted value. Padding elements were excluded from the reconstruction loss. The KL divergence term is

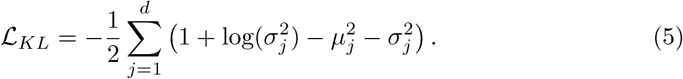

We used the MAE as a metric to represent the algorithm’s performance. The MAE was calculated over the same non-padding distance-matrix elements used for the reconstruction loss. It is worth mentioning that the distance matrices are supposed to be symmetric. We perform the symmetrization of the predicted distance matrices as a part of postprocessing as follows:

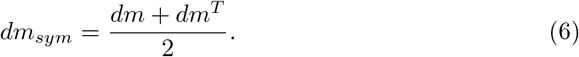

If symmetrization is performed before calculating the loss function, training becomes unstable because a set of values for the predicted distance matrix could satisfy the same value for the symmetric distance matrix. Hence, the sum would matter for the specific element of the symmetric matrix, making the values more flexible by adding an extra degree of freedom, and resulting in unstable training. Therefore, symmetrization was not included in the training loss and was applied as a postprocessing step to the predicted distance matrices. Metrics were calculated for both the original and symmetrized predictions.

After recovering the three-dimensional point coordinates using multidimensional scaling (MDS), we obtained a spatial representation of the RNA structure. We used classical MDS, which reconstructs coordinates through eigendecomposition of the double-centered squared-distance matrix. The mathematical formulation is described below.

The full inference pipeline comprised the following steps:

- Prediction of the inter-nucleotide centroid distance matrix by the VAE from the RNA sequence and stochastic input.
- Symmetrization of the predicted distance matrix and reconstruction of three-
- dimensional nucleotide centroid coordinates using multidimensional scaling (MDS).
- Template-based assembly of the atomic RNA structure using the reconstructed nucleotide centroid coordinates.
- Addition of missing atoms required to obtain a complete atomic model.
- Molecular-dynamics refinement of the reconstructed structure to reduce steric clashes and unfavorable local geometries.

After predicting the inter-nucleotide distance matrix, three-dimensional nucleotide coordinates were reconstructed using MDS and used as target coordinates for template-based atomic reconstruction. The template library was constructed from RNA structures in the PDB by extracting consecutive four-nucleotide fragments and grouping them according to nucleotide sequence. This procedure produced 256 sequence-specific template catalogs containing a total of 46,199 fragments.

For reconstruction, overlapping consecutive four-nucleotide windows were generated from the target RNA sequence with a step size of one nucleotide. For each window, candidate templates from the corresponding sequence-specific catalog were aligned to the reconstructed nucleotide coordinates, and the template with the lowest RMSD to the predicted points was selected. The selected template was then transformed into the coordinate frame of the corresponding predicted nucleotide positions.

Because consecutive four-nucleotide windows overlap, atoms of a given nucleotide could receive coordinate estimates from more than one aligned template. When multiple estimates were available for the same atom, the final Cartesian coordinates were obtained by averaging the corresponding coordinate estimates across the contributing templates.

The reconstructed RNA structures were refined using molecular dynamics (MD) simulations in OpenMM [25]. The refinement implementation selected the first force-field combination compatible with the reconstructed RNA structure from a predefined sequence of AMBER-based parameter sets, beginning with AMBER14 RNA.OL3 combined with TIP3P-FB water, followed by AMBER14 RNA.OL3 with TIP3P and additional AMBER fallback parameterizations. The RNA was solvated in an explicit water box with 2.0 nm padding and an ionic strength of 0.15 M using Na^+^ and Cl^−^ ions. Long-range electrostatic interactions were treated using the particle-mesh Ewald (PME) method with a 1.0 nm nonbonded cutoff, and bonds involving hydrogen atoms were constrained. Simulations were performed at 300 K using a Langevin-middle integrator with a friction coefficient of 1 ps^−1^ and a timestep of 0.02 fs.

Before MD refinement, the system was energy-minimized. Positional restraints were applied to RNA heavy atoms and progressively reduced over 50 refinement stages, with the restraint force constant logarithmically decreasing from 10^2^ to 10^−1^ kJ mol^−1^ nm^−2^. Each stage consisted of 5,000 MD steps followed by energy minimization. After the restrained stages, an additional 5,000 MD steps were performed without positional restraints, followed by a final energy minimization. The resulting RNA coordinates were then extracted and saved as the refined structure.

Because distance-based reconstruction is invariant under reflection, MDS does not determine absolute chirality. No explicit chirality-selection step is used here; therefore, a reconstructed structure may in principle be reflected. Addressing this ambiguity explicitly is left for future work.

Figure 5 illustrates the full AutoRNA reconstruction workflow. Starting from the FASTA sequence, the VAE predicts structural priors that are converted into spatial constraints and transformed into a coarse 3D model via distance geometry, which is then matched to the closest structural template. The final structure is refined using molecular dynamics to obtain an MD-refined reconstructed RNA structure, resulting in the final PDB structure.

**Fig. 5.**
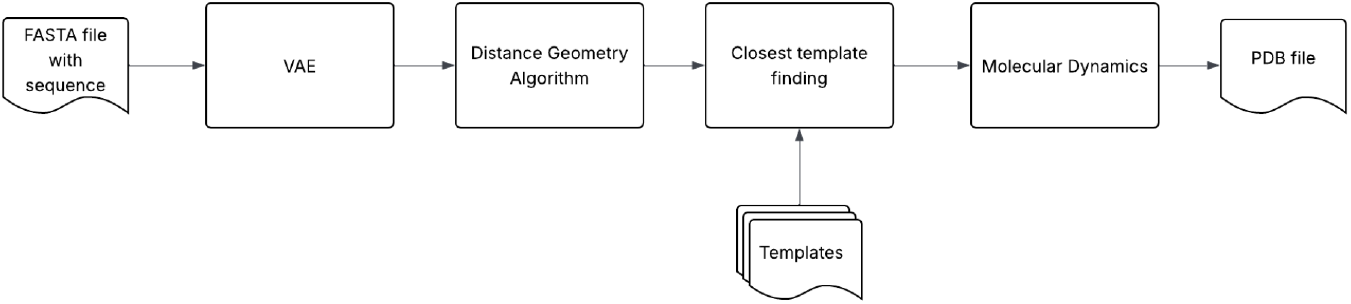
Full AutoRNA reconstruction workflow.

## 3 Results

For the experiment, we trained our model for 1000 epochs. We used the validation subset for fine-tuning the parameters, and the testing was only for assessing the final prediction. We stored the model with the best validation MAE. For training, the batch size was set to 16, which was chosen to obtain a stable solution while achieving a sufficiently fast learning rate. However, we did not observe a significant correlation between batch size and final performance. We also used the Adam optimizer [26]. For the first experiment, the minimum and maximum lengths of the RNA sequences were 4 and 64 nucleotides, respectively. We used the best-performing loss parameter, which balances the KL loss and reconstruction loss, namely that which equals 1.1. The training time was approximately 2.1 hours (Apple M1 processor). The inference time for one RNA sequence during batch execution was approximately 0.001 seconds.

In the inference stage, we also assembled the final PDB file containing the final structure from a set of templates. The most suitable template was selected based on its proximity to the predicted points. Next, we optimized the assembled structure using molecular dynamics methods. As our experiments show, the nucleotide centroids of the structure obtained in this way differ from the points obtained from VAE by no more than 1 Å; thus, they do not affect the final results. In addition to RMSE and MAE, we calculated the global distance test (GDT) and template modeling (TM) score after symmetrization. We calculated the GDT score based on the distances between the corresponding nucleotide centroids in the predicted and reference structures, evaluated at multiple distance thresholds. We used the values of 1, 2, 4, and 8 Å, according to [27]. We calculated the TM score based on [28] using the tmscoring Python library [29]. We included the GDT and TM scores in the final computation due to the limitations of the RMSE (and MAE) metric for different nucleotide lengths.

Table 1 presents the results of the experiment for the training, validation, and testing subsets. The results (RMSE, MAE, RMSE + sym, and MAE + sym) were slightly better for the testing subset than for the validation subset due to the stochasticity of the values. Interestingly, the GDT score was significantly lower for the testing subset than for the training and validation subsets. For the testing subset, we achieved an RMSE of 4.49 Åand an MAE of 3.13 Å. The difference between training and validation performance was consistent with some degree of overfitting, suggesting that additional training data could improve model generalization. Note that we also report performance after symmetrization of the predicted distance matrices during postprocessing. Symmetrization had only a negligible effect on the reported error metrics. For the testing subset, the RMSE changed from 4.49 Å before symmetrization to 4.48 Å after symmetrization, while the corresponding MAE changed from 3.13 Å to 3.12 Å. Thus, enforcing the expected symmetry of the distance matrix produced only a very small change in predictive error. The symmetrized results are reported to quantify the effect of this postprocessing step rather than as evidence of improved model performance. It is worth mentioning that the test set was small (37 sequences), and the result that the performance on the test set is slightly better than on validation is possible due to high variance on the test set.

**Table 1.** The results of the variational autoencoder (VAE) experiment for the training, validation, and testing subsets.

|  | MAE | MAE+sym | RMSE | RMSE+sym | GDT-score | TM-score |
| --- | --- | --- | --- | --- | --- | --- |
| Train | $3.23 \pm 1.64$ | $3.22 \pm 1.64$ | $4.69 \pm 2.46$ | $4.69 \pm 2.46$ | $39.67 \pm 20.58$ | $0.25 \pm 0.14$ |
| Validation | $3.86 \pm 2.61$ | $3.86 \pm 2.61$ | $5.56 \pm 3.77$ | $5.56 \pm 3.77$ | $26.83 \pm 20.87$ | $0.18 \pm 0.05$ |
| Test | $3.13 \pm 1.69$ | $3.12 \pm 1.69$ | $4.49 \pm 2.42$ | $4.48 \pm 2.43$ | $32.13 \pm 20.19$ | $0.19 \pm 0.07$ |

Figure 6 presents the predicted and reference distance matrices for two representative RNA sequences. We employed the distance geometry method to reconstruct the coordinates from the distance matrix, a classic technique proven effective in nuclear magnetic resonance spectroscopy [30]. The challenge is to identify a set of coordinates that optimally fits the given distance constraints. According to [31], this problem can be approached by reformulating it as the task of finding a matrix:

**Fig. 6.**
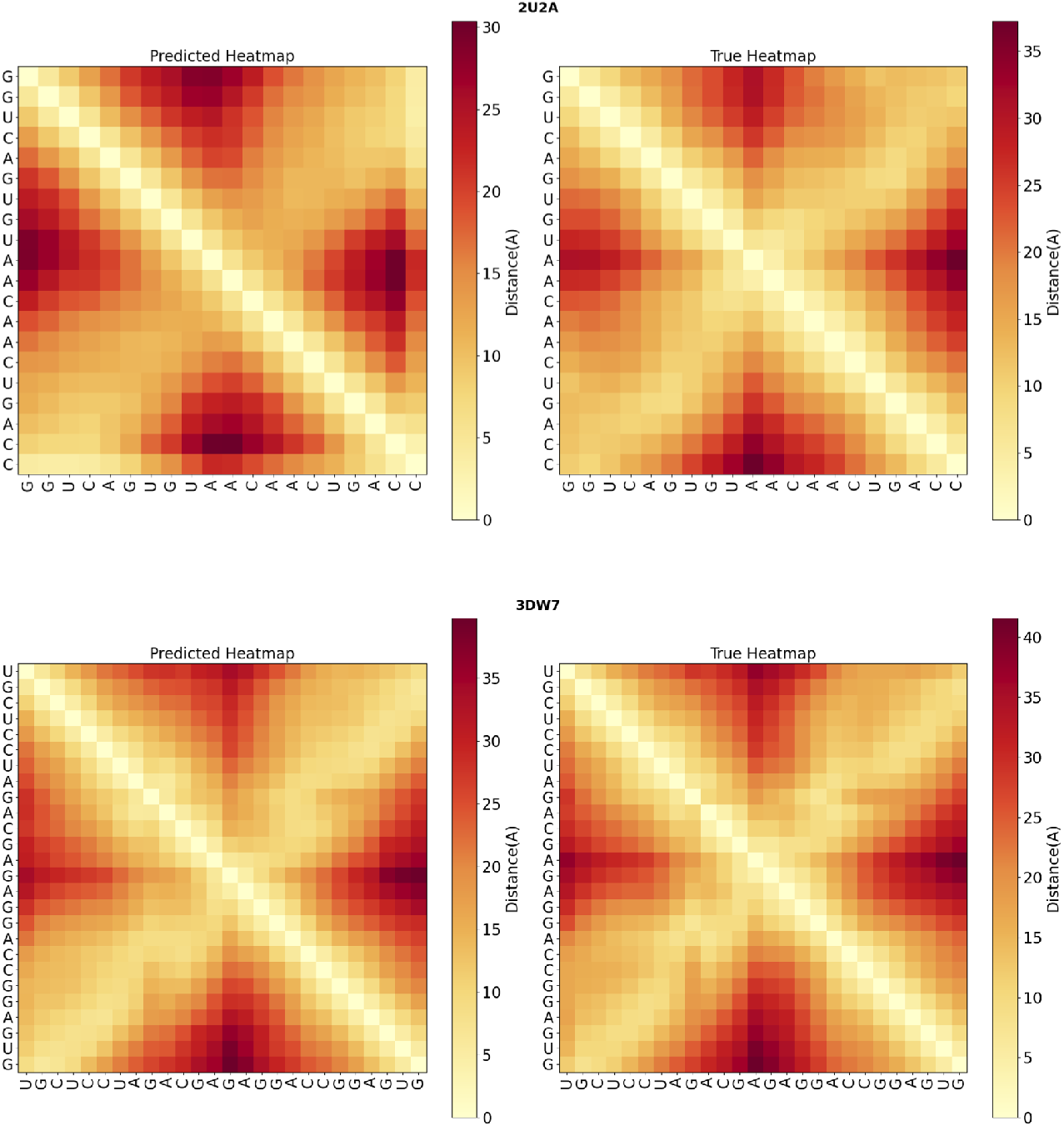
Predicted (left) and reference (right) inter-nucleotide distance matrices for two representative RNA structures: stem-loop IIa from *Saccharomyces cerevisiae* U2 snRNA (PDB: 2U2A, top) and the sarcin/ricin domain from *Escherichia coli* 23S rRNA (PDB: 3DW7, bottom).

Let *D*^(2)^ denote the matrix of squared pairwise distances, with elements 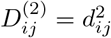. Classical multidimensional scaling first constructs the centered Gram matrix

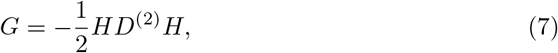

where

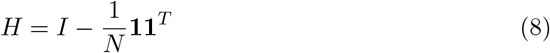

is the centering matrix. The Gram matrix is then eigendecomposed as

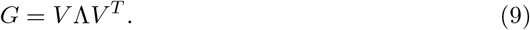

The three-dimensional Cartesian coordinates are recovered from the three largest positive eigenvalues and their corresponding eigenvectors:

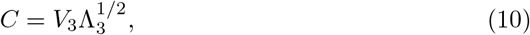

where *V*_3_ contains the eigenvectors corresponding to the three largest positive eigenvalues, Λ_3_ is the diagonal matrix containing these eigenvalues, and *C* ∈ ℝ^*N ×*3^ contains the reconstructed three-dimensional coordinates of the nucleotide centroids. In addition to the basic experiments, we examined the sensitivity of the predicted distance matrices to the stochastic input. For several RNA sequences, we generated distance matrices five times using different noise vectors, as illustrated in Figure 7. Variation in the noise vector produced relatively small changes in the resulting predictions. Across the test set, the average coefficient of variation of the prediction error across different noise realizations was

**Fig. 7.**
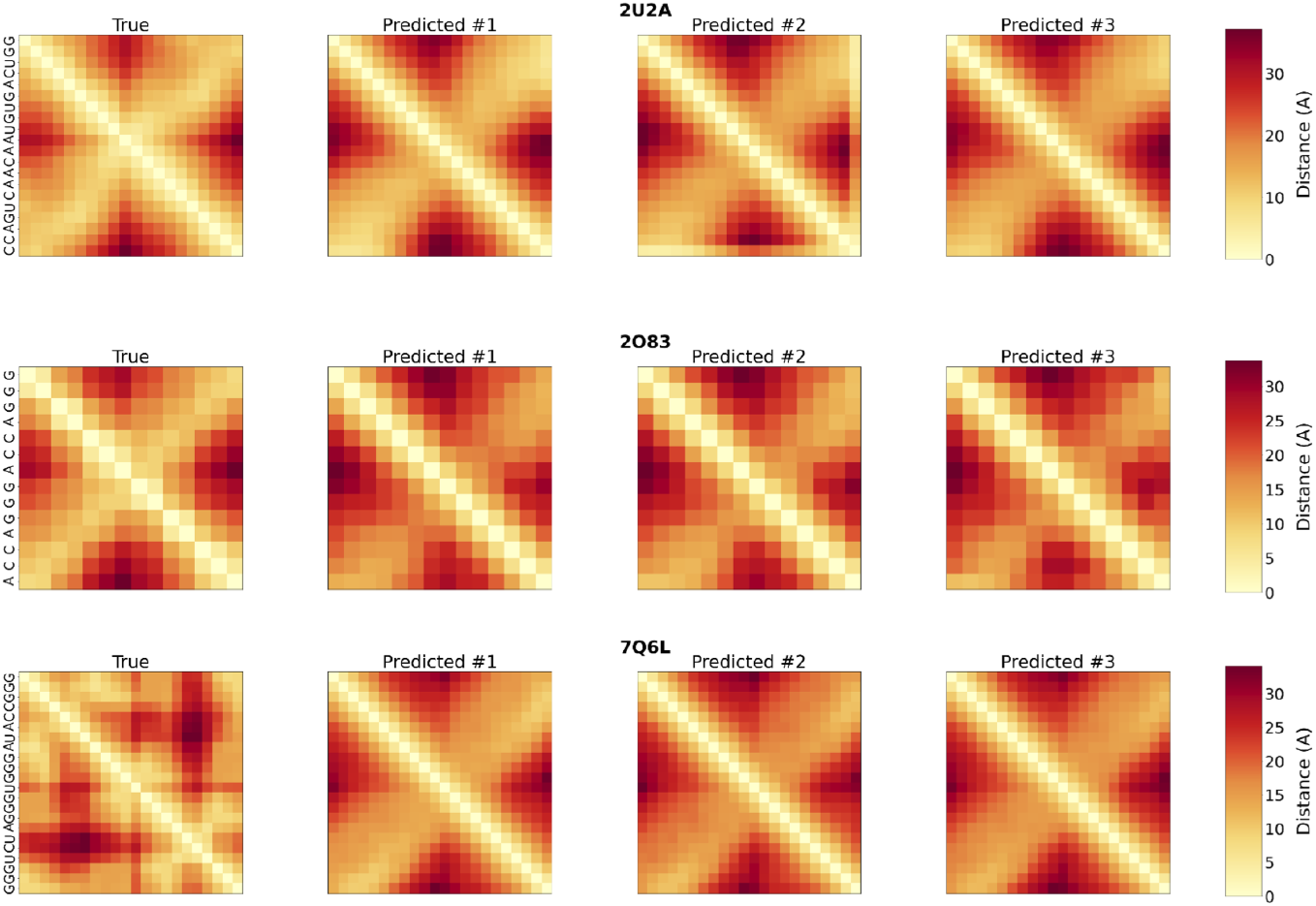
Structures generated with different noise vectors (three noise vectors for each sequence) for three RNA structures (from top to bottom: 2U2A, 2O83, 7Q6L). The image on the left represents the true distance matrices, and the images on the right represent the predicted structures with different initial noise vectors. The nucleotide labels corresponding to the sequences are shown on the left.

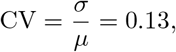

where *µ* and *σ* denote the mean and standard deviation, respectively, of the prediction error across noise realizations. Thus, under the evaluated sampling procedure, the prediction error showed relatively low variability with respect to changes in the noise vector. We report this result descriptively and do not infer the organization or information content of the latent representation from this analysis alone.

For each predicted distance matrix, we quantified deviations from Euclidean embeddability using the negative eigen-fraction (NEF). Let *λ*_1_, …, *λ*_*N*_ denote the eigenvalues of the centered Gram matrix *G*. We define NEF as

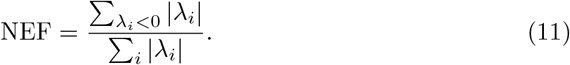

A value of zero indicates that the centered Gram matrix has no negative eigenvalues, whereas positive values quantify the relative magnitude of its negative spectral components. On the test set, the mean NEF was 0.092. Thus, approximately 9.2% of the total absolute eigenvalue magnitude was associated with negative eigenvalues under this definition. This indicates that the predicted distance matrices contain measurable non-Euclidean components. Because no validated threshold was used to classify a predicted matrix as Euclidean or non-Euclidean, the NEF value is reported here as a descriptive measure of deviation from Euclidean embeddability rather than as evidence that the predicted matrices satisfy Euclidean criteria.

In the next experiment, we trained the same model but for longer RNA sequences, fixing the maximum length at 128 nucleotides while keeping the other parameters unchanged. The total dataset in this case consisted of 925 sequences, compared to 852 sequences containing at most 64 nucleotides. As the training, validation, and testing subsets naturally differed from those when the maximum length was fixed at 64 nucleotides, we compared the results with those in Table 1, considering the stochastic component. Table 2 presents the results of the experiments in the testing subset. The decrease in accuracy for longer RNA sequences may reflect several factors, including the smaller number of available long-RNA structures, increased structural complexity and longer-range dependencies, and limitations associated with applying the same fixed model architecture to longer sequences. The present experiments do not distinguish the relative contributions of these factors. To assess overfitting quantitatively, we compared performance across the training, validation, and held-out test subsets. The validation MAE (3.86 Å) and RMSE (5.56 Å) were moderately higher than the corresponding training values (3.23 Åand 4.69 Å), representing an approximate 18–20% increase in error on unseen data. Structural quality metrics exhibited a similar trend: the mean GDT score decreased from 39.7 to 26.8 and the TM-score from 0.25 to 0.18 between training and validation subsets, indicating reduced global structural alignment. The test subset yielded performance comparable to the validation subset (MAE = 3.13 Å, RMSE = 4.49 Å). However, because dataset partitioning was based on sequence similarity rather than structural similarity, these results should not be interpreted as demonstrating generalization to structurally independent RNA molecules.

**Table 2.** Comparison of the performance of the AutoRNA algorithm for small and medium-sized RNA sequences in the testing subset.

| Test dataset | Number of sequences | RMSE | GDT-score |
| --- | --- | --- | --- |
| Sequences, $\leq 64$ nt | 30 | $4.49 \pm 2.35$ | $32.13 \pm 20.19$ |
| Sequences, 65–128 nt | 7 | $6.05 \pm 1.17$ | $14.72 \pm 4.33$ |

**Table 3.** Comparison of mean absolute error (MAE) for AutoRNA, RhoFold, and SimRNA on the test dataset, stratified by RNA sequence length.

|  | AutoRNA | RhoFold | SimRNA |
| --- | --- | --- | --- |
| MAE ( Å), $\leq 32$ nts | 3.475 | 3.414 | 3.990 |
| MAE ( Å), $> 32$ nts | 4.248 | 3.692 | 5.125 |

We then compared AutoRNA with several representative contemporary methods for 3D RNA structure prediction: AlphaFold3 [4], Boltz-2 [32], SimRNA [48], and RhoFold [33]. These methods were selected to represent complementary computational strategies relevant to RNA structure prediction. RhoFold was included as an RNA-focused deep-learning method for tertiary structure prediction, AlphaFold3 and Boltz-2 as general deep-learning models for biomolecular structure prediction, and SimRNA as an RNA-specific coarse-grained conformational-sampling method based on a knowledge-based statistical potential. For simulations with RhoFold, we used standard parameter settings (10000 iterations). The compared methods use substantially different structure-prediction and refinement strategies. AutoRNA includes an optional molecular-dynamics refinement stage, whereas AlphaFold3, Boltz-2, and RhoFold primarily rely on learned structure-prediction pipelines. SimRNA instead uses a coarse-grained representation, conformational sampling, and a knowledge-based statistical potential. To distinguish the contribution of the learned AutoRNA prediction from that of subsequent physical refinement, we report AutoRNA results both with and without molecular dynamics.

For modeling with AlphaFold3 and Boltz-2, we used the AF3 server from Google and the Boltz-2-NIM server from Nvidia, respectively. We selected single-chain structures shorter than 64 nucleotides from the RNA-Puzzles competition [5], identifying nine structures that satisfied these criteria. Because of the small number of eligible targets, this analysis should be regarded as a limited illustrative external benchmark rather than a comprehensive evaluation of performance across RNA-Puzzles targets. Only one RNA (puzzle 16) in our dataset was shorter than 32 nucleotides. The RNA sequences had different origins: they were catalytic RNAs (puzzle 16), riboswitches (puzzles 29 and 38), and aptamers (puzzle 32). The root mean square deviation (RMSD) metric was used for comparison. We used PyMOL [34] to calculate the RMSD between the experimental and modeled RNA structures, and the results for each structure are presented in Table 4. The PDB files with the structures and the FASTA file with the sequences are provided in the Supplementary Materials.

**Table 4.**
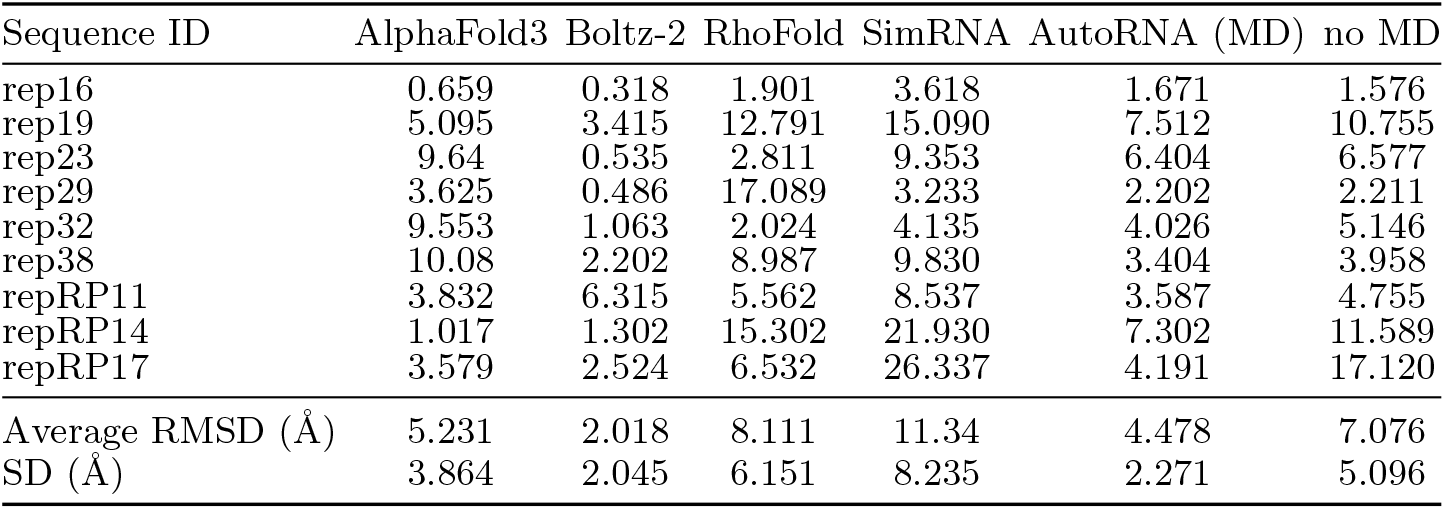
Comparison of the algorithms (AlphaFold3, Boltz-2, RhoFold, SimRNA) with our algorithm (AutoRNA) on the subset of RNA-puzzles dataset.

| Sequence ID | AlphaFold3 | Boltz-2 | RhoFold | SimRNA | AutoRNA (MD) | no MD |
| --- | --- | --- | --- | --- | --- | --- |
| rep16 | 0.659 | 0.318 | 1.901 | 3.618 | 1.671 | 1.576 |
| rep19 | 5.095 | 3.415 | 12.791 | 15.090 | 7.512 | 10.755 |
| rep23 | 9.64 | 0.535 | 2.811 | 9.353 | 6.404 | 6.577 |
| rep29 | 3.625 | 0.486 | 17.089 | 3.233 | 2.202 | 2.211 |
| rep32 | 9.553 | 1.063 | 2.024 | 4.135 | 4.026 | 5.146 |
| rep38 | 10.08 | 2.202 | 8.987 | 9.830 | 3.404 | 3.958 |
| repRP11 | 3.832 | 6.315 | 5.562 | 8.537 | 3.587 | 4.755 |
| repRP14 | 1.017 | 1.302 | 15.302 | 21.930 | 7.302 | 11.589 |
| repRP17 | 3.579 | 2.524 | 6.532 | 26.337 | 4.191 | 17.120 |
| Average RMSD (Å) | 5.231 | 2.018 | 8.111 | 11.34 | 4.478 | 7.076 |
| SD (Å) | 3.864 | 2.045 | 6.151 | 8.235 | 2.271 | 5.096 |

Overall, MD refinement reduced the mean RMSD, although the magnitude and direction of the effect varied across individual structures. It should be noted that AutoRNA predictions are reported both with and without molecular dynamics (MD) refinement, whereas models such as AlphaFold3 do not incorporate physics-based post-processing as part of their inference pipeline. Therefore, AutoRNA results without MD are provided for a more direct comparison with purely data-driven methods. While the enhancement was modest in most cases, three initially poorly predicted structures showed a substantial improvement of approximately 3–4 Å after MD refinement. These results indicate that the effect of MD refinement varied across structures, with larger improvements observed for some structures with higher initial prediction errors. The present analysis does not establish the mechanism underlying these differences. Although the positions of individual nucleotides may differ between the predicted and experimental structures, the modeled structures retain some aspects of the overall backbone organization. Figure 8 illustrates representative comparisons between the predicted and experimental RNA structures.

**Fig. 8.**
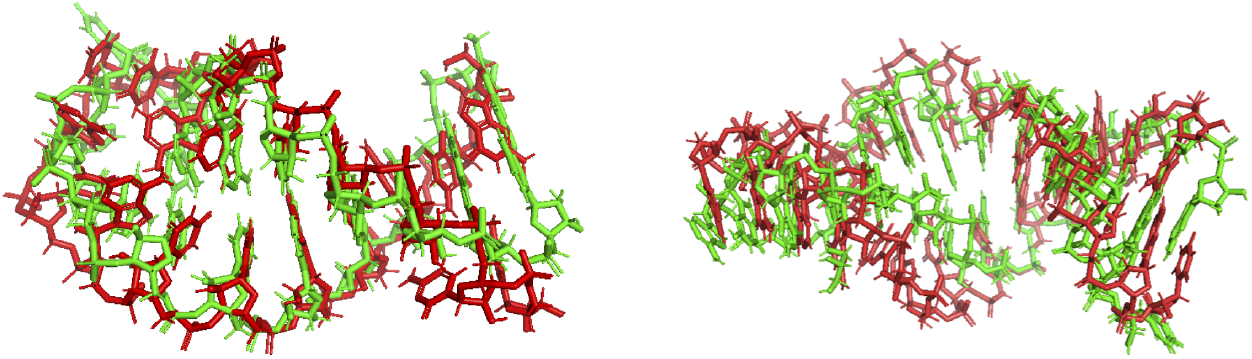
Comparison of the experimental and predicted structures for RNA stem-loops 6XXB (left) and 7L3R (right).

Additional examples comparing predicted and experimental RNA structures are provided in Appendix A.

## 4 Discussion

The proposed AutoRNA approach has several limitations despite its relatively simple formulation. The main drawback is that the predicted and learning set distance matrices have a fixed and limited size. Thus, the PDB dataset contains insufficient long-RNA structures for proper training. Another limitation arises from the imbalance between the number of trainable parameters in the VAE and the size of the available RNA structural dataset. Although the model contains approximately 64.5 million parameters, dropout (rate = 0.5) was used to reduce the risk of overfitting, while sequence-similarity-based clustering was used during dataset partitioning to reduce sequence redundancy between the training, validation, and test subsets. Because this partitioning was based on sequence similarity rather than structural similarity, it does not guarantee structural independence between the subsets, and structurally related RNAs may therefore occur across different subsets.

Although the current performance of AutoRNA (average RMSE ≈ 4.5, Å, TM-score *<* 0.25) does not indicate near-native tertiary-structure recovery, the predicted distance matrices capture coarse-grained structural information. AutoRNA should therefore be interpreted primarily as a generator of sequence-conditioned coarse-grained structural restraints rather than as a standalone tertiary-structure predictor. Such restraints could potentially be incorporated into downstream reconstruction or physics-based refinement workflows. Related hybrid strategies have been used in RNA structure-prediction frameworks, where learned structural restraints are combined with external reconstruction, optimization, or refinement procedures [15, 16].

An additional experiment on RNA sequences containing 65–128 nucleotides showed a decline in performance relative to the primary experiment on shorter sequences. This decline may reflect several factors, including the smaller number of available long-RNA structures, increased structural complexity and longer-range dependencies, and limitations associated with applying the same fixed model architecture to longer sequences. The present experiments do not distinguish the relative contributions of these factors. Therefore, the primary analysis was restricted to RNA sequences containing at most 64 nucleotides, for which substantially more structural data were available. The architecture of the VAE is one of the possible approaches for resolving RNA folding. Additionally, incorporating RNA-specific features into the model based on the biophysical properties of RNA molecules could be attempted. For example, we could consider the secondary structure of RNA [35] or the functional class of predicted RNA molecules or features obtained from RNA templates as additional features.

Another potential direction is the computational characterization of long non-coding RNAs, whose structural and interaction properties remain challenging to characterize [41, 42]. However, extending AutoRNA to long non-coding RNAs would require substantially improved performance on longer RNA sequences and considerably larger and more diverse structural training datasets. In addition, chemical modifications can influence RNA structure and function, suggesting that nucleotide-modification information could be considered as an additional feature in future extensions of the model [47].

Evolutionary information can provide valuable constraints for RNA structure modeling. Covariation analysis of multiple-sequence alignments is well established for identifying evolutionarily conserved RNA base pairs and secondary-structure features [20, 49, 50]. Comparative RNA-analysis methods can therefore exploit evolutionary information when sufficiently informative sequence alignments are available. However, covariation evidence for conserved base pairing should not be interpreted as directly specifying a complete three-dimensional RNA structure, and deriving detailed tertiary structural constraints from evolutionary information remains more challenging. In addition, the statistical power of covariation analysis depends on the depth, diversity, and evolutionary variation of the available sequence alignment. Incorporating evolutionary features into AutoRNA therefore represents a potential direction for future work, particularly for RNA families with sufficiently informative alignments. Future extensions could also incorporate physically informed neural-network components that explicitly account for molecular constraints and atomic interactions.

Another important direction is to enhance the dataset. Modeling additional RNA structures using physics-based methods, such as molecular dynamics or Monte Carlo sampling, could provide candidate structures for dataset augmentation, although generating and validating such structures would require additional computational resources and careful quality control. Thus, using a rotamer library may be a practical solution, as it would allow changing one or several nucleotides in an RNA molecule into other nucleotides. Their local conformation must be in accordance with the rotamer library, after which several molecular dynamics steps could be performed to remove clashes and refine the structure.

## 5 Conclusions

This study developed AutoRNA, a VAE-based approach for learning sequence-conditioned coarse-grained structural representations of RNA molecules. Unlike computational approaches based primarily on molecular simulation [11, 25], AutoRNA learns nonlinear relationships between RNA sequence and inter-nucleotide spatial organization directly from structural data. The resulting predictions should be interpreted as coarse-grained structural priors rather than near-native tertiary structures and may provide initial structural information for downstream reconstruction and physics-based refinement.

In conclusion, AutoRNA provides a proof-of-concept demonstration that a sequence-conditioned variational autoencoder can learn coarse-grained inter-nucleotide distance representations from a relatively small dataset of experimentally determined RNA structures. On the test dataset, the model achieved an RMSE of approximately 4.5 Å for predicted inter-nucleotide distances. However, the modest GDT and TM-scores indicate that these predictions should not be interpreted as near-native tertiary structures. The present results therefore support AutoRNA primarily as an approach for learning coarse-grained structural restraints under limited-data conditions. Future work with larger and more structurally diverse datasets, improved structural reconstruction, and more rigorous evaluation of generalization will be required to determine the utility of such representations for RNA structure modeling.

Nonetheless, the AutoRNA algorithm has limitations regarding the specific RNA sequences because of the need for appropriate training data. For some RNA structures, sub-optimal conformations that could vary from the original tertiary structures were found. However, the proposed AutoRNA algorithm may be used to generate sequence-conditioned structural priors that support downstream tertiary structure modeling workflows.

## Acknowledgements

The authors thank A. Kasianov, A. Krasnaya, S. Semyonov, and A. Safaraleev for valuable discussions and support.

## Declarations

The following sections provide the required declarations in accordance with journal submission guidelines.

### Ethics approval and consent to participate

Not applicable.

### Consent for publication

Not applicable.

### Data availability

The datasets generated and/or analyzed during the current study are publicly available in the Zenodo repository: https://doi.org/10.5281/zenodo.16887142 The dataset used in this study was derived from publicly available macromolecular structure records deposited in the Protein Data Bank (PDB; wwPDB, https://www.wwpdb.org/). Specifically, we selected PDB entries containing RNA-only structures according to the filtering criteria described in the Methods section.

### Materials availability

Not applicable.

### Code availability

The code used for data preprocessing, model training, inference, three-dimensional structure reconstruction, and molecular-dynamics refinement is publicly available in the AutoRNA GitHub repository (https://github.com/quantori/AutoRNA). The repository also provides the software dependencies and configuration parameters required to reproduce the computational workflow. The trained model checkpoint used for the reported experiments is provided together with the associated model configuration and random-seed settings.

### Author contributions

Conceptualization: M.K., L.U., Y.G.; Methodology: M.K., L.U.; Data curation: M.K., L.U., F.Z.; Software: M.K.; Formal analysis: M.K.; Investigation: M.K.; Visualization: M.K., F.Z., I.P.; 3D structure reconstruction: L.U.; Writing – original draft: M.K., L.U.; Writing – review & editing: M.K., F.Z., I.P., O.K.; Supervision: Y.G.; Project administration: Y.G., O.K. All authors have read and approved the final manuscript.

## Appendix A Additional structural comparisons

Green color reflects the actual structure; red color reflects the predicted structure.

**1ESH** ↓

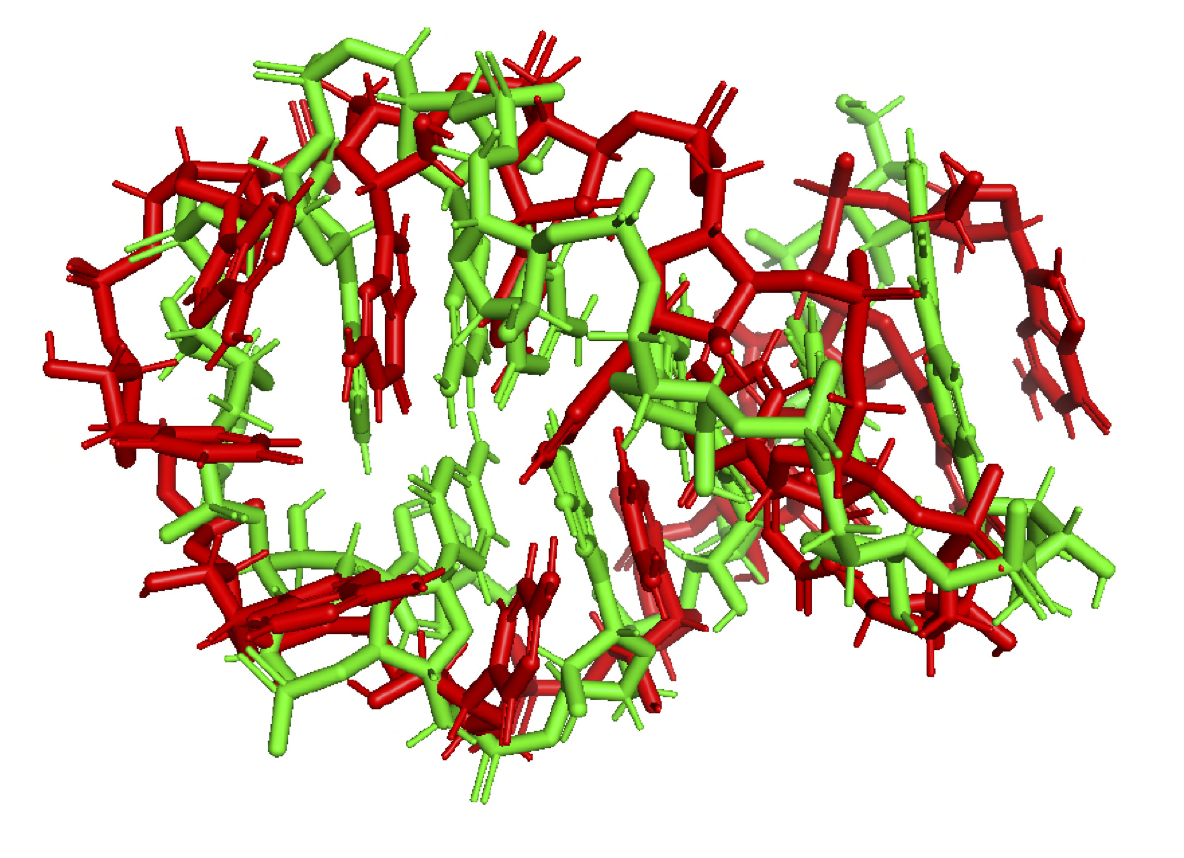

**2G1G** ↓

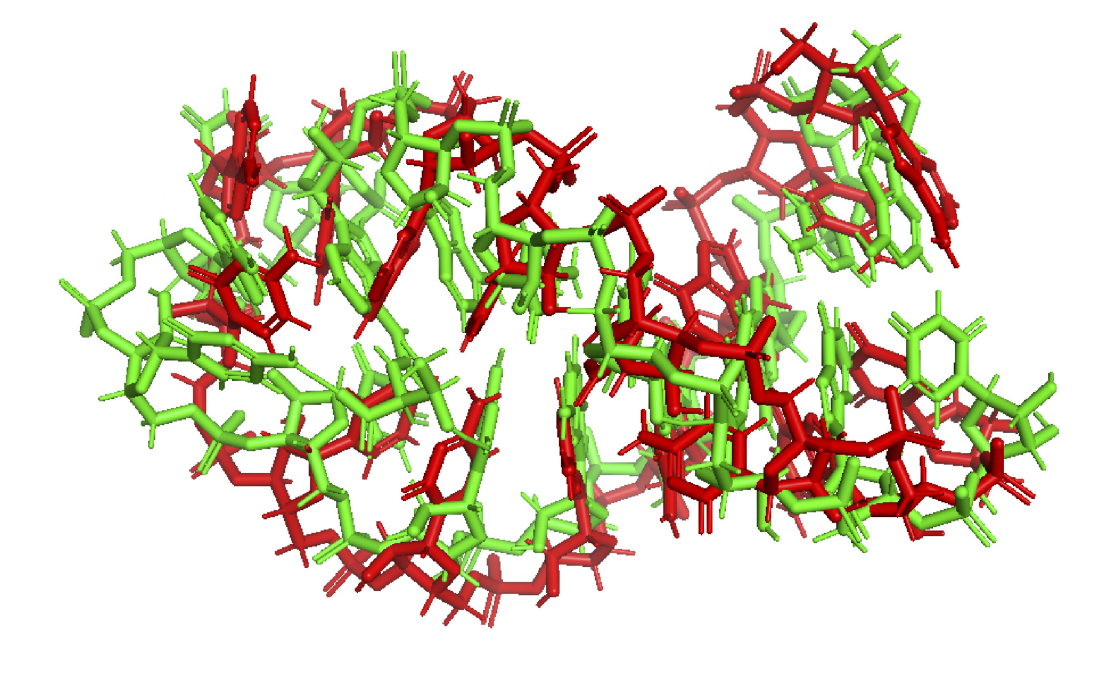

**1I46** ↓

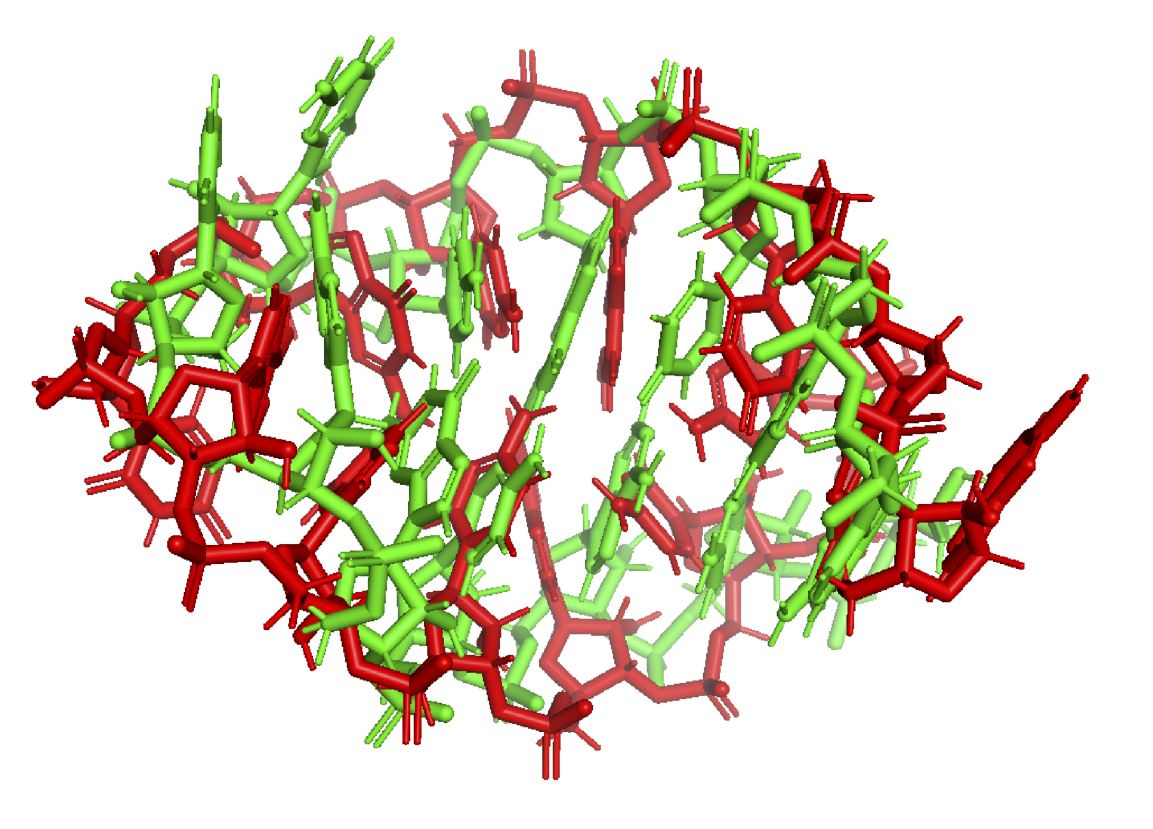

**1ROQ** ↓

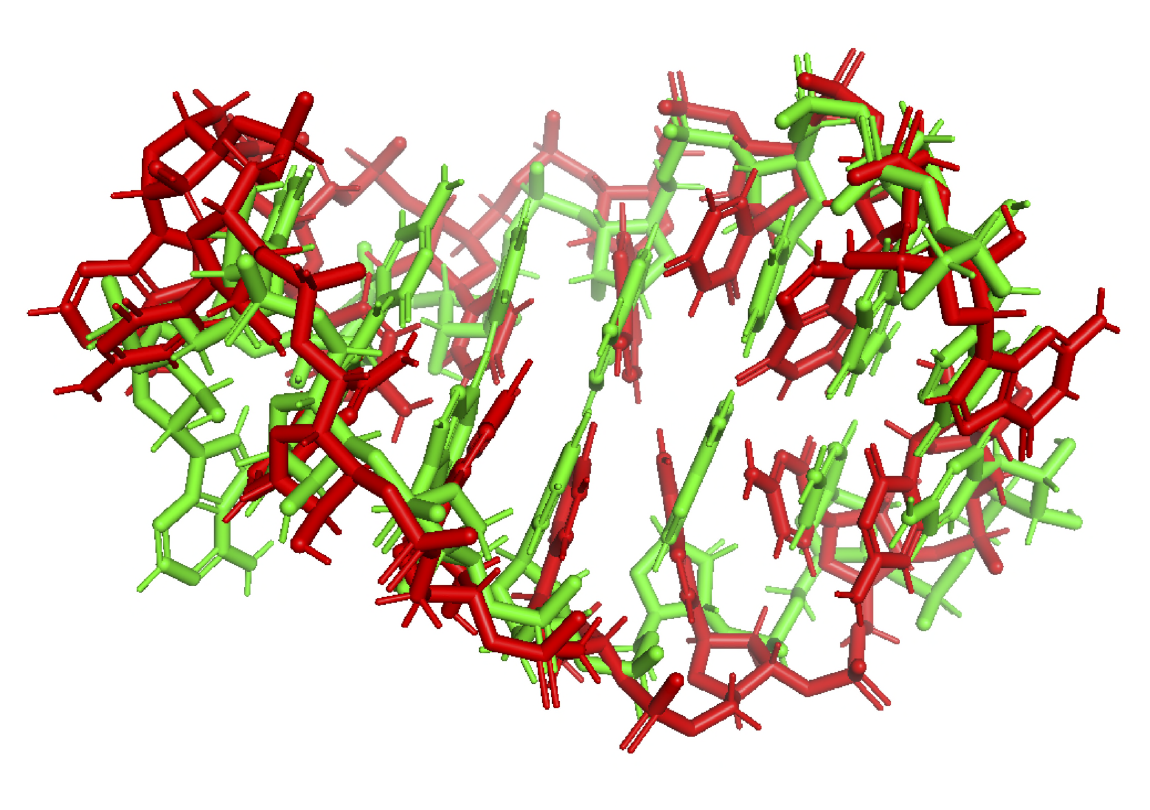

**1NZ1** ↓

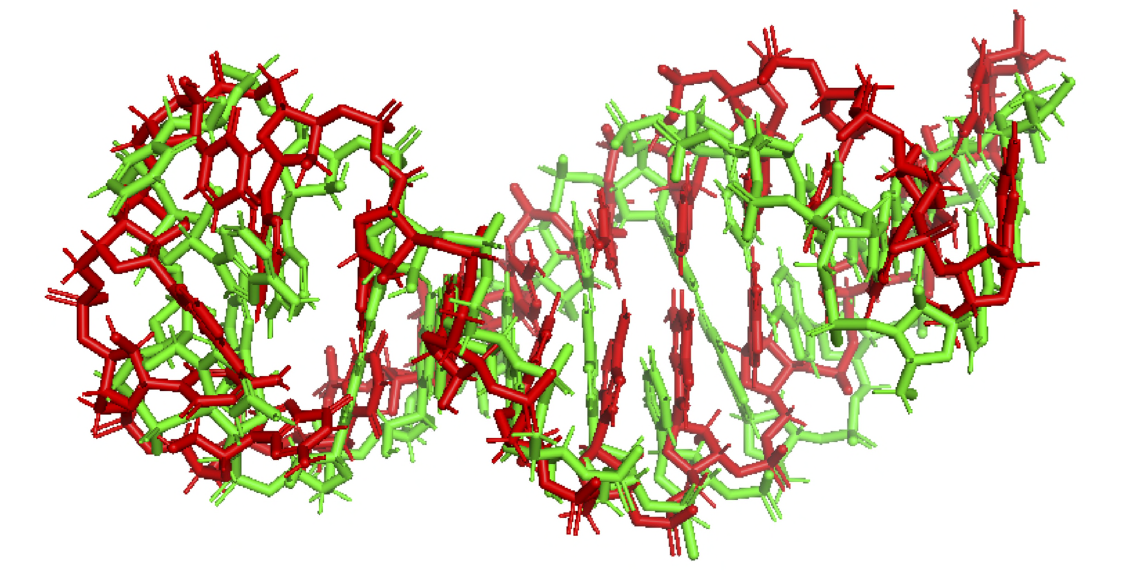

**2LPA** ↓

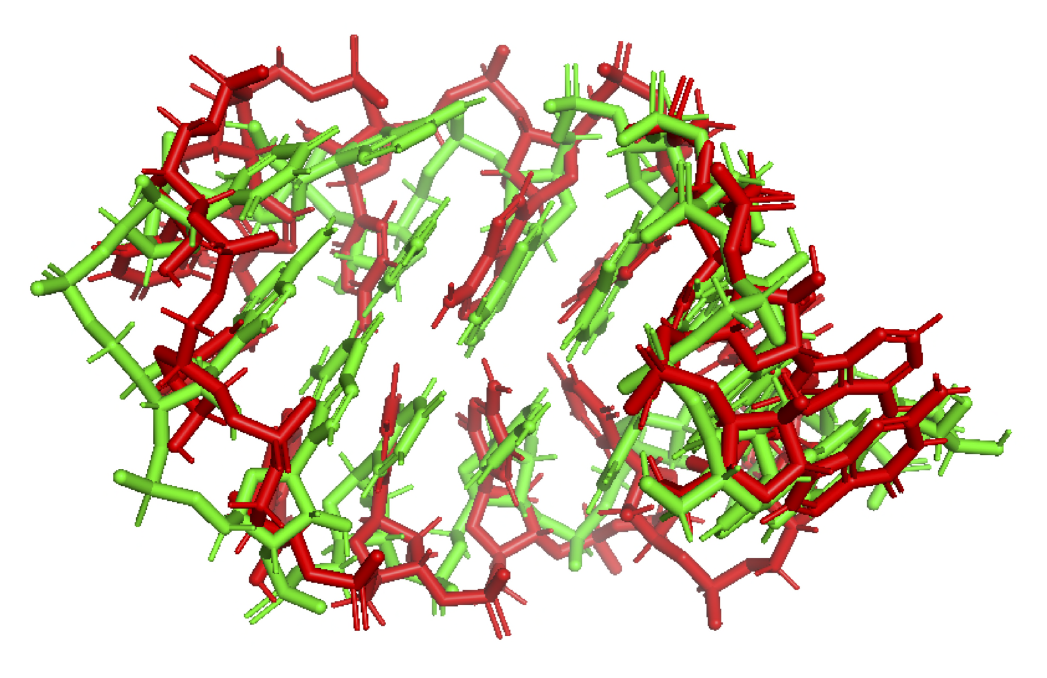

## References

[1] Miao Z, Westhof E. RNA Structure: Advances and Assessment of 3D Structure Prediction. Annu Rev Biophys. 2017;46:483–503. doi:10.1146/annurev-biophys-070816-034125.

[2] Cao X, Zhang Y, Ding Y, et al. Identification of RNA structures and their roles in RNA functions. Nat Rev Mol Cell Biol. 2024;25:784–801.

[3] Jumper, J. et al. Highly accurate protein structure prediction with AlphaFold. Nature 596(7873), 583–589 (2021)

[4] Abramson, J. et al. Accurate structure prediction of biomolecular interactions with AlphaFold 3. Nature 630(8016), 493–500 (2024). doi:10.1038/s41586-024-07487-w

[5] Cruz, J. A. et al. RNA-Puzzles: a CASP-like evaluation of RNA three-dimensional structure prediction. RNA 18(4), 610–625 (2012)

[6] Danaee, P. et al. bpRNA: large-scale automated annotation and analysis of RNA secondary structure. Nucleic Acids Res. 46(11), 5381–5394 (2018)

[7] Gillespie, J., Mayne, M., Jiang, M. RNA folding on the 3D triangular lattice. BMC Bioinformatics 10, 369 (2009). doi:10.1186/1471-2105-10-369

[8] Sharma, S., Ding, F., Dokholyan, N. V. iFoldRNA: three-dimensional RNA structure prediction and folding. Bioinformatics 24(17), 1951–1952 (2008)

[9] Tan, Y. L. et al. rsRNASP: a residue-separation-based statistical potential for RNA 3D structure evaluation. Biophys. J. 121(1), 142–156 (2022)

[10] Rahmanov, S., Kulakovskiy, I., Uroshlev, L., Makeev, V. Empirical potentials for ion binding in proteins. J. Bioinform. Comput. Biol. 8(3), 427–435 (2010)

[11] Case, D. A. et al. The Amber biomolecular simulation programs. J. Comput. Chem. 26(16), 1668–1688 (2005)

[12] Brooks, B. R. et al. CHARMM: the biomolecular simulation program. J. Comput. Chem. 30(10), 1545–1614 (2009)

[13] Al-Hashimi, H. M., Walter, N. G. RNA dynamics: it is about time. Curr. Opin. Struct. Biol. 18(3), 321–329 (2008)

[14] Li, J., Chen, S. J. RNA 3D structure prediction using coarse-grained models. Front. Mol. Biosci. 8, 720937 (2021)

[15] Wang, W. et al. trRosettaRNA: automated prediction of RNA 3D structure with transformer network. Nat. Commun. 14(1), 7266 (2023)

[16] Watkins, A. M., Rangan, R., Das, R. FARFAR2: improved de novo Rosetta prediction of complex global RNA folds. Structure 28(8), 963–976.e6 (2020). doi:10.1016/j.str.2020.05.011

[17] Fu, L. et al. UFold: fast and accurate RNA secondary structure prediction with deep learning. Nucleic Acids Res. 50(3), e14 (2022)

[18] Burley, S. K. et al. Protein Data Bank (PDB): the single global macromolecular structure archive. Methods Mol. Biol. 1607, 627–641 (2017)

[19] Hong, X. et al. RR3DD: an RNA global structure-based RNA three-dimensional structural classification database. RNA Biol. 18(Suppl 2), 738–746 (2021). doi:10.1080/15476286.2021.1989200

[20] Rivas, E. RNA structure prediction using positive and negative evolutionary information. PLoS Comput. Biol. 16(10), e1008387 (2020)

[21] Li, W., Godzik, A. Cd-hit: clustering large sets of sequences. Bioinformatics 22(13), 1658–1659 (2006)

[22] Doersch, C. Tutorial on variational autoencoders. arXiv:1606.05908 (2016)

[23] Asperti, A., Trentin, M. Balancing Reconstruction Error and Kullback-Leibler Divergence in Variational Autoencoders. IEEE Access 8, 199440–199448 (2020). doi:10.1109/ACCESS.2020.3034828

[24] Burgess, C. P. et al. Understanding disentangling in beta-VAE. arXiv:1804.03599 (2018)

[25] Eastman, P. et al. OpenMM 7: Rapid development of high performance algorithms for molecular dynamics. PLoS Comput. Biol. 13(7), e1005659 (2017). doi:10.1371/journal.pcbi.1005659

[26] Kingma, D. P., Ba, J. Adam: a method for stochastic optimization. arXiv:1412.6980 (2014)

[27] Zemla, A. LGA: a method for finding 3D similarities in protein structures. Nucleic Acids Res. 31(13), 3370–3374 (2003)

[28] Zhang, Y., Skolnick, J. TM-align: a protein structure alignment algorithm based on the TM-score. Nucleic Acids Res. 33(7), 2302–2309 (2005). doi:10.1093/nar/gki524

[29] Menéndez Hurtado, D. tmscoring: Python implementation of the TMscore program, version 0.4.post0 (2019). Available at: https://pypi.org/project/tmscoring/

[30] Mucherino, A., Lavor, C., Liberti, L., Maculan, N. (eds.). Distance Geometry: Theory, Methods, and Applications. Springer, New York (2013). doi:10.1007/978-1-4614-5128-0

[31] Trosset, M. W. A new formulation of the nonmetric strain problem in multidimensional scaling. J. Classif. 15(1), 15–35 (1998).

[32] Passaro, S. et al. Boltz-2: Towards Accurate and Efficient Binding Affinity Prediction. bioRxiv, 2025.06.14.659707 (2025). doi:10.1101/2025.06.14.659707

[33] Shen, T. et al. Accurate RNA 3D structure prediction using a language model-based deep learning approach. Nat. Methods 21(12), 2287–2298 (2024). doi:10.1038/s41592-024-02487-0

[34] DeLano, W. L. PyMOL molecular graphics system. Available at: https://pymol.org (2002)

[35] Vicens, Q., Kieft, J. S. Thoughts on how to think (and talk) about RNA structure. Proc. Natl. Acad. Sci. USA 119(17), e2112677119 (2022). doi:10.1073/pnas.2112677119

[36] Qiao, Y. et al. DeepFusion: A deep bimodal information fusion network for unraveling protein-RNA interactions using in vivo RNA structures. Comput. Struct. Biotechnol. J. 23, 617–625 (2024). doi:10.1016/j.csbj.2023.12.040

[37] Florentino, B. R. et al. BioPrediction-RPI: Democratizing the prediction of interaction between non-coding RNA and protein with end-to-end machine learning. Comput. Struct. Biotechnol. J. 23, 2267–2276 (2024). doi:10.1016/j.csbj.2024.05.031

[38] Krautwurst, S., Lamkiewicz, K. RNA-protein interaction prediction without high-throughput data: An overview and benchmark of in silico tools. Comput. Struct. Biotechnol. J. 23, 4036–4046 (2024). doi:10.1016/j.csbj.2024.11.015

[39] Sun, L. Z. et al. RLDOCK: A New Method for Predicting RNA–Ligand Interactions. J. Chem. Theory Comput. 16(11), 7173–7183 (2020). doi:10.1021/acs.jctc.0c00798

[40] Pan, X. et al. RBPsuite: RNA-protein binding sites prediction suite based on deep learning. BMC Genomics 21, 884 (2020). doi:10.1186/s12864-020-07291-6

[41] Cicconetti, C. et al. 3plex enables deep computational investigation of triplex forming lncRNAs. Comput. Struct. Biotechnol. J. 21, 3091–3102 (2023). doi:10.1016/j.csbj.2023.05.016

[42] Ballarino, M. et al. Exploring the landscape of tools and resources for the analysis of long non-coding RNAs. Comput. Struct. Biotechnol. J. 21, 4706–4716 (2023). doi:10.1016/j.csbj.2023.09.041

[43] Cheetham, S. W., Gruhl, F., Mattick, J. S., Dinger, M. E. Long noncoding RNAs and the genetics of cancer. Br. J. Cancer 108(12), 2419–2425 (2013). doi:10.1038/bjc.2013.233

[44] Gupta, R. A. et al. Long non-coding RNA HOTAIR reprograms chromatin state to promote cancer metastasis. Nature 464(7291), 1071–1076 (2010). doi:10.1038/nature08975

[45] Braga, E. A. et al. Ten Hypermethylated lncRNA Genes Are Specifically Involved in the Initiation, Progression, and Lymphatic and Peritoneal Metastasis of Epithelial Ovarian Cancer. Int. J. Mol. Sci. 25(21), 11843 (2024). doi:10.3390/ijms252111843

[46] Brovkina, O. I. et al. Identification of Novel Differentially Expressing Long Non-Coding RNAs with Oncogenic Potential. Mol. Biol. 55(4), 548–554 (2021). doi:10.1134/S0026893321020175

[47] Pichot, F. et al. Quantification of substoichiometric modification reveals global tsRNA hypomodification, preferences for angiogenin-mediated tRNA cleavage, and idiosyncratic epitranscriptomes of human neuronal cell-lines. Comput. Struct. Biotechnol. J. 21, 401–417 (2023). doi:10.1016/j.csbj.2022.12.020

[48] Moafinejad, S. N. et al. SimRNAweb v2.0: a web server for RNA folding simulations and 3D structure modeling, with optional restraints and enhanced analysis of folding trajectories. Nucleic Acids Res. 52(W1), W368–W373 (2024). doi:10.1093/nar/gkae356

[49] Rivas, E., Clements, J., Eddy, S. R. A statistical test for conserved RNA structure shows lack of evidence for structure in lncRNAs. Nat. Methods 14, 45–48 (2017). doi:10.1038/nmeth.4066

[50] Rivas, E. RNA covariation at helix-level resolution for the identification of evolutionarily conserved RNA structure. PLoS Comput. Biol. 19, e1011262 (2023). doi:10.1371/journal.pcbi.1011262

